## Supplementary Information for "Structure and dynamics of cholesterol-mediated aquaporin-0 arrays and implications for lipid rafts"

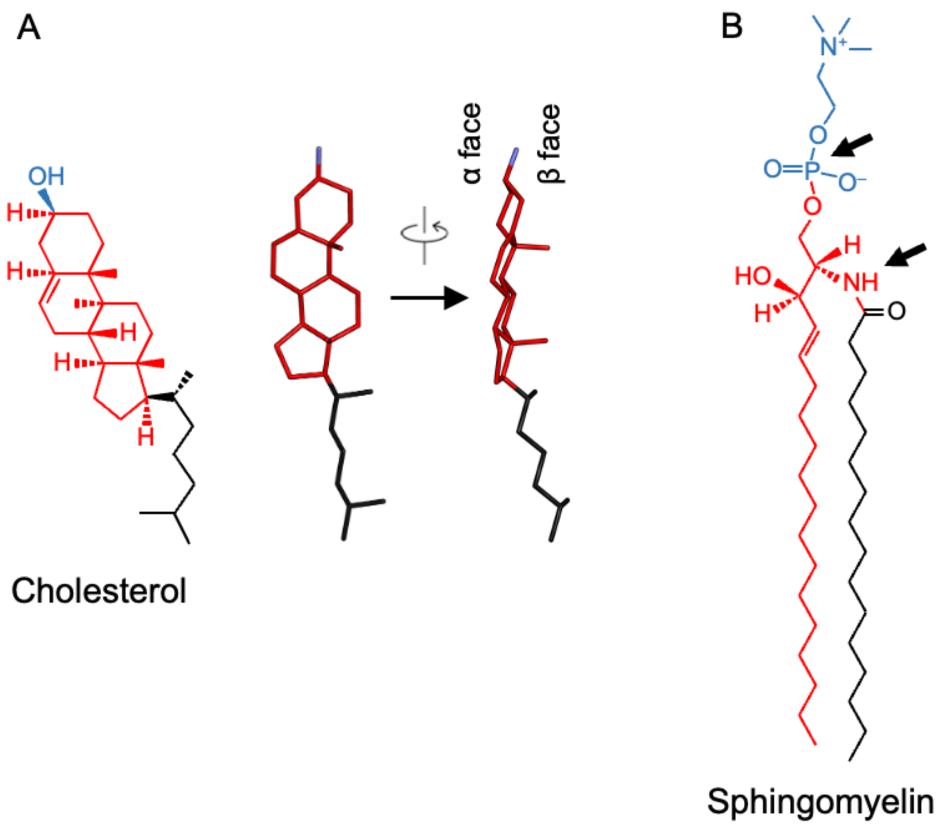


**Figure 1–figure supplement 1.** Chemical structures of raft lipids. (**A**) Cholesterol consists of a planar four-aromatic ring structure (red) with a short isooctyl alkyl chain (black) and a small 3-β-hydroxyl head group (blue). The side view shows the asymmetric distribution of the aliphatic groups linked to the ring system and the resulting smooth (α) and rough (β) faces of the cholesterol molecule. (**B**) Sphingomyelin consists of a sphingosine amino alcohol (red) with a fatty acid (black) and phosphocholine head group (blue).

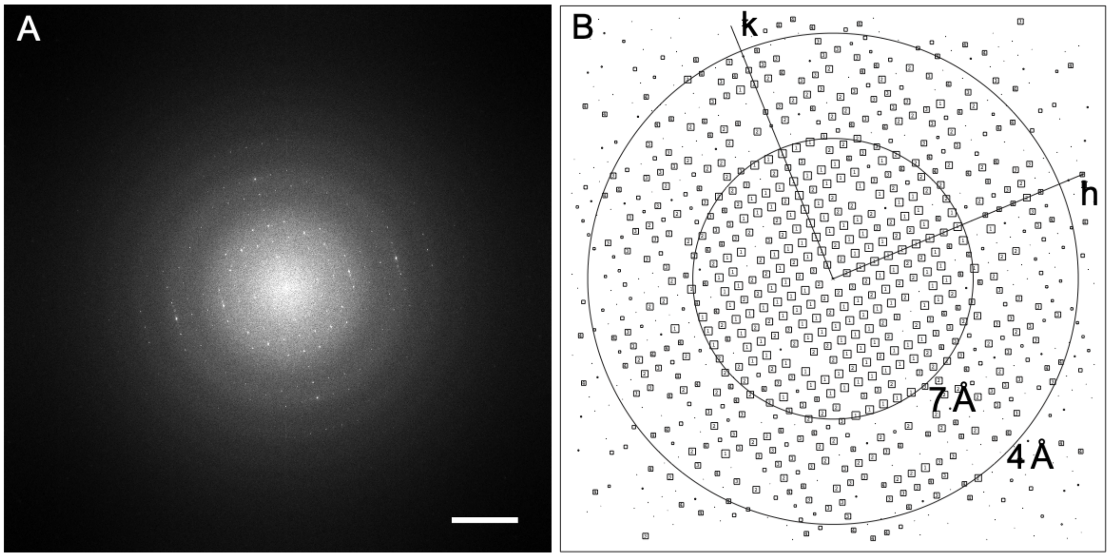


**Figure 1–figure supplement 2.** Diffraction of an image of an AQP0 2D crystal grown with pure cholesterol. (**A**) Power spectrum of a raw image of an AQP0 2D crystal in a pure cholesterol membrane. The scale bar represents (10 Å)$^{-1}$. (**B**) Intensity quotient (IQ) plot after computational lattice unbending. After lattice unbending of the image in *2dx* (Gipson et al., 2007b), the IQ plot showed reflections with IQ values of 3 (corresponding to a peak-to-background ratio of 2.3) to a resolution better than 4 Å.

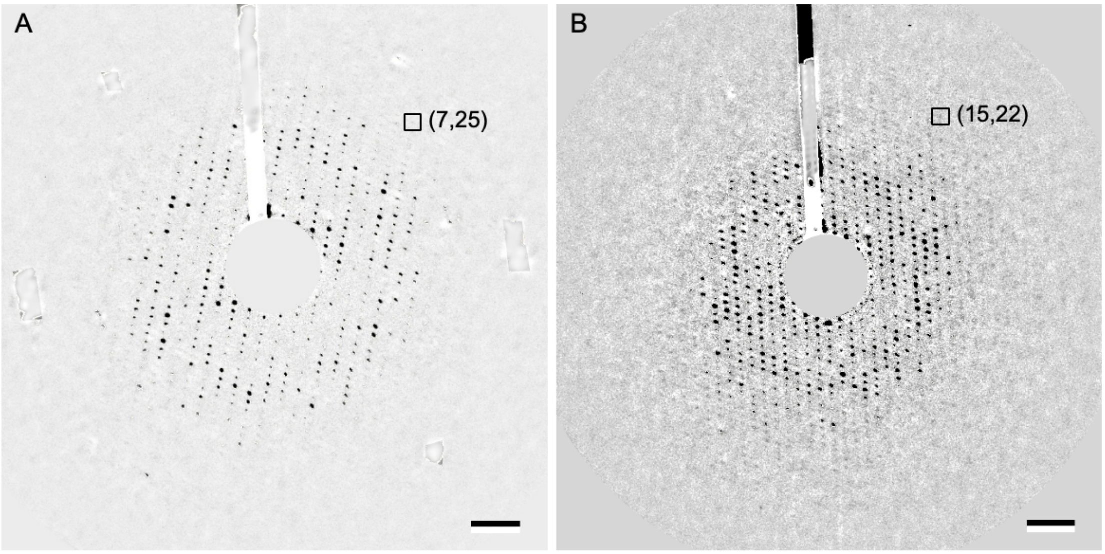


**Figure 2–figure supplement 1.** Electron diffraction patterns of 2D crystals reconstituted with sphingomyelin/cholesterol mixtures tilted to 60º. (**A**) Background-subtracted electron diffraction pattern of a 60º tilted AQP0 2D crystal reconstituted at a molar sphingomyelin:cholesterol ratio of 2:1. The boxed reflection corresponds to a resolution of 2.52 Å resolution. (**B**) Background-subtracted electron diffraction pattern of a 60º tilted AQP0 2D crystal reconstituted at a molar sphingomyelin:cholesterol ratio of 1:2. The boxed reflection corresponds to a resolution of 2.46 Å resolution. Scale bars indicate (10 Å)$^{-1}$.

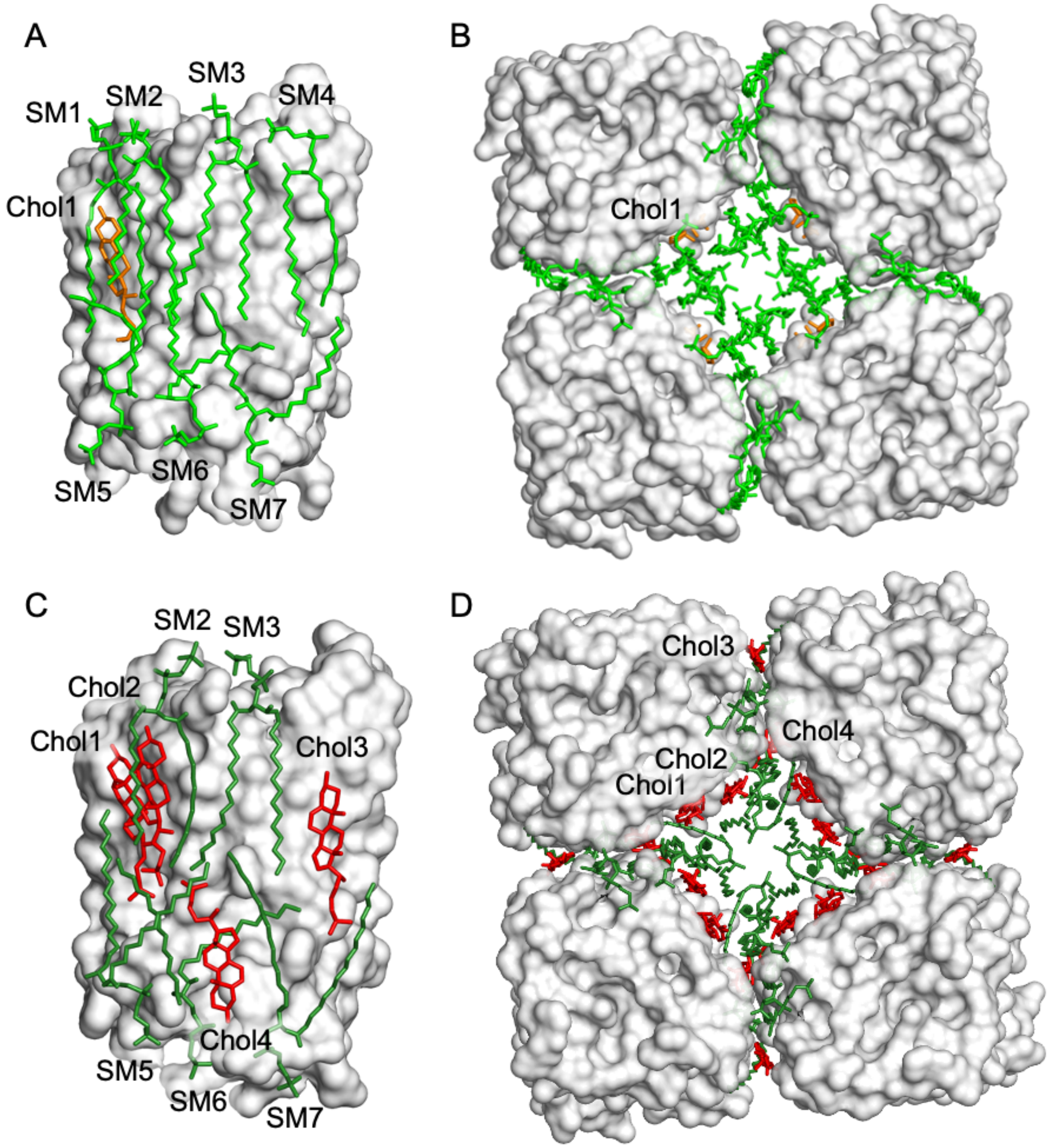


**Figure 3–figure supplement 1.** Distribution of sphingomyelin and cholesterol molecules around AQP0 in 2D crystals. (**A–B**) 2D crystals grown with a molar sphingomyelin:cholesterol ratio of 2:1. (**C–D**) 2D crystals grown with a molar sphingomyelin:cholesterol ratio of 1:2**.** Lipid distribution in AQP0 arrays formed with a molar sphingomyelin:cholesterol ratio of 2:1. Panels (A) and (C) show the side view of a single AQP0 subunit, and panels (B) and (D) show a top view of four subunits (from different tetramers) in the area where four different tetramers come together in the array. Note that lipids at the crystallographic four-fold axis could not be modeled.

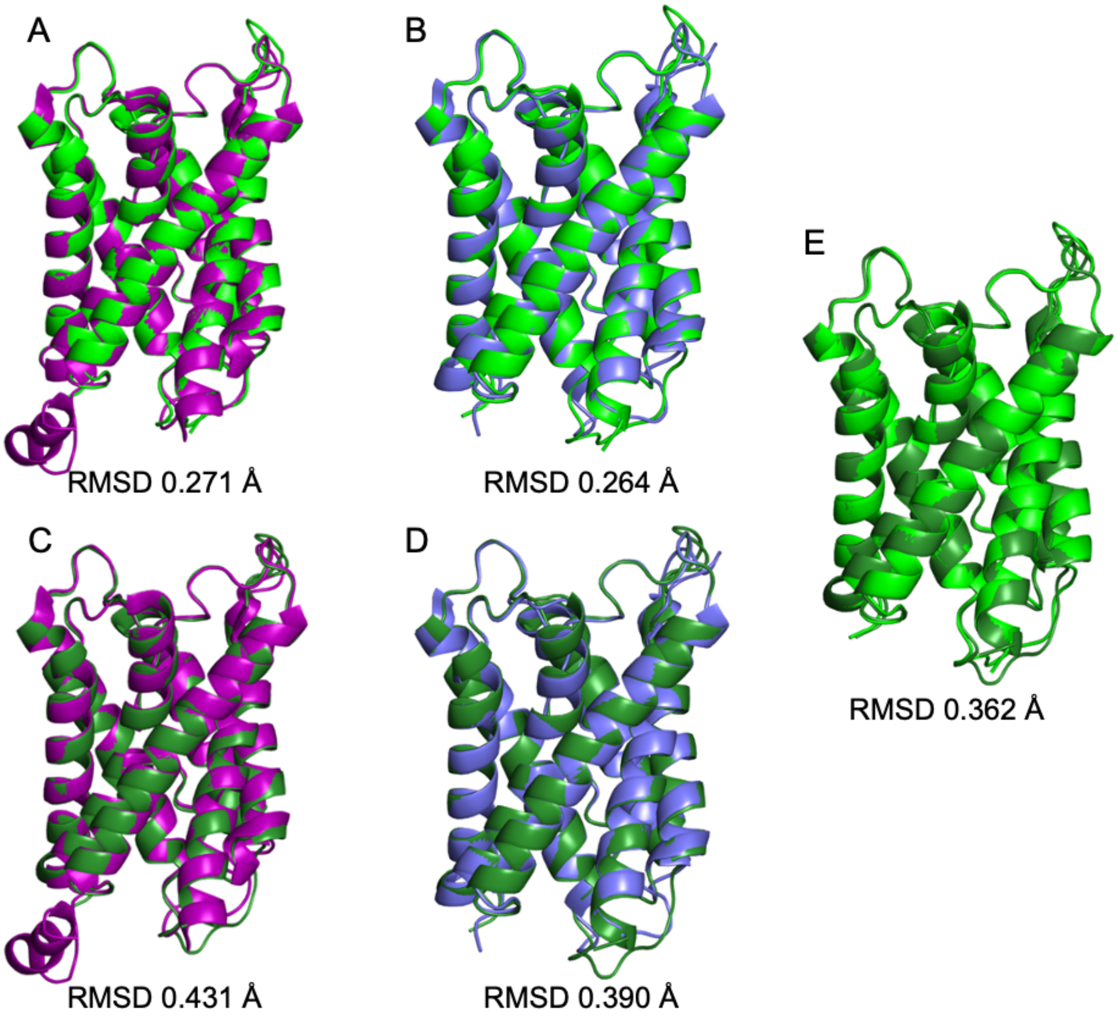


**Figure 3–figure supplement 2.** Structure comparisons of AQP0 in different lipid bilayers. The root mean square deviation (RMSD) between the Cα atoms of AQP0 structures determined in membranes formed with different lipids is very low and variations between structures are largely constrained to extramembranous loops. (**A**) $AQP0_{2SM:1Chol}$ (light green) *versus* $AQP0_{DMPC}$ (purple). (**B**) $AQP0_{2SM:1Chol}$ (light green) *versus* $AQP0_{EPL}$ (blue). (**C**) $AQP0_{1SM:2Chol}$ (dark green) *versus* $AQP0_{DMPC}$ (purple). (**D**) $AQP0_{1SM:2Chol}$ (dark green) *versus* $AQP0_{DMPC}$ (purple). (**E**) $AQP0_{2SM:1Chol}$ (light green) *versus* $AQP0_{1SM:2Chol}$ (dark green).

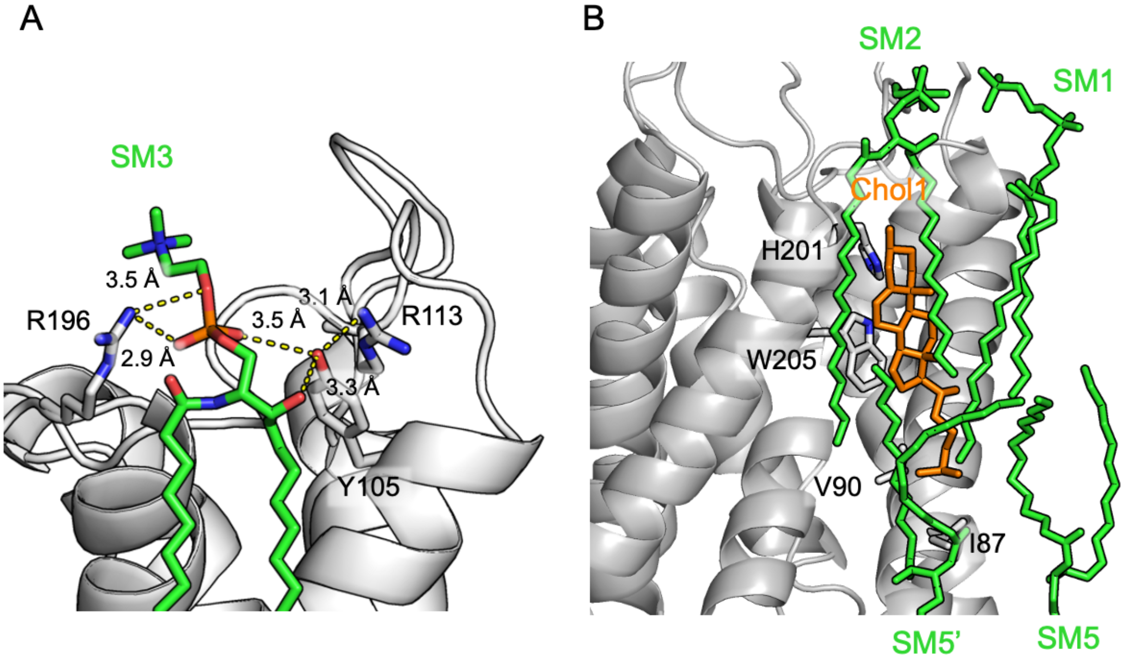


**Figure 3–figure supplement 3.** Interactions of sphingomyelin lipids in $AQP0_{2SM:1Chol}$ with AQP0 and Chol1. (**A**) The head group of sphingomyelin SM3 interacts with AQP0 residues Tyr-105 and Arg-196. Yellow dash lines indicate the hydrogen bonds. (**B**) The acyl chains of sphingomyelins SM1, SM2, SM5, and SM5' have hydrophobic interactions with cholesterol Chol1, as well as AQP0 residues of His-201, Trp-205, Val-90, and Ile-87.

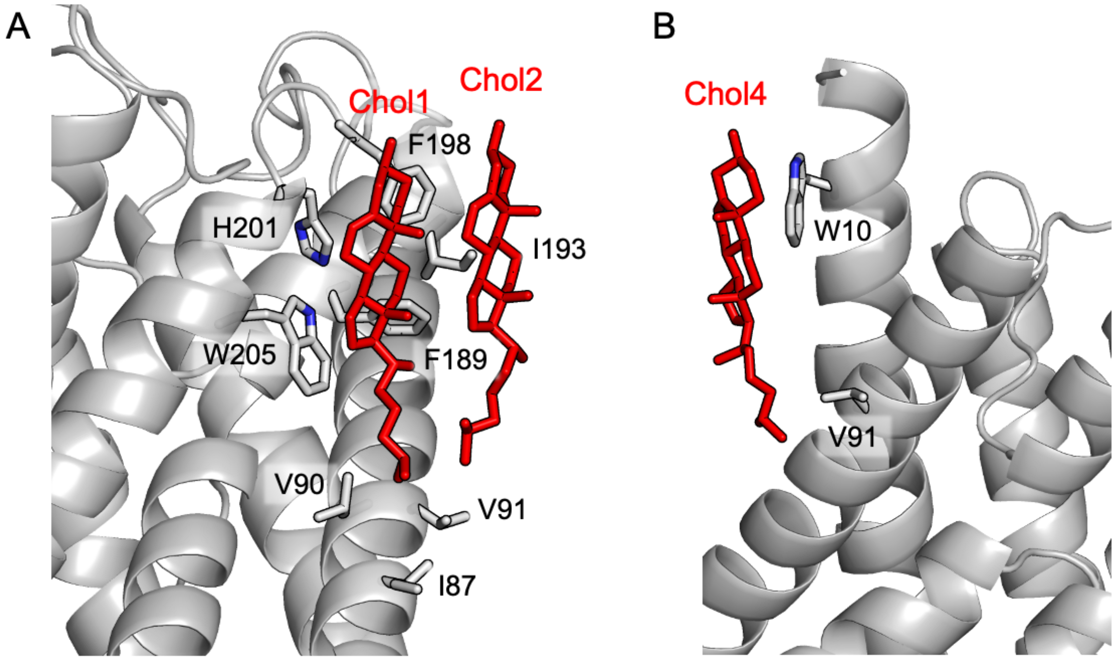


**Figure 4–figure supplement 1.** Interactions of cholesterol molecules in $AQP0_{1SM:2Chol}$ with AQP0. (**A**) The cholesterols in the extracellular leaflet, Chol1 and Chol2. (**B**) The cholesterol in the cytoplasmic leaflet, Chol4. All cholesterol molecules interact with hydrophobic residues on the AQP0 surface. The ring systems of Chol1, Chol2 and Chol4 also make π-stacking interactions with aromatic residues Trp-205, Phe-198 and Trp-10, respectively.

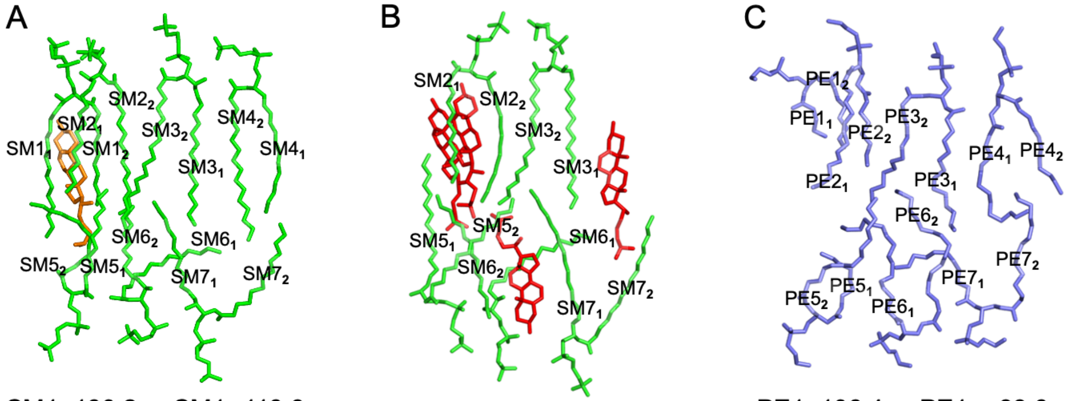


| | | |
|---|---|---|
| SM$1_1$ 130.2 ; SM$1_2$ 113.9 | | PE$1_1$ 106.4 ; PE$1_2$ 99.3 |
| SM$2_1$ 117.7 ; SM$2_2$ 84.1 | SM$2_1$ 78.7 ; SM$2_2$ 91.6 | PE$2_1$ 69.7 ; PE$2_2$ 74.4 |
| SM$3_1$ 72.0 ; SM$3_2$ 70.9 | SM$3_1$ 59.7 ; SM$3_2$ 65.7 | PE$3_1$ 50.0 ; PE$3_2$ 53.8 |
| SM$4_1$ 92.2 ; SM$4_2$ 85.0 | | PE$4_1$ 64.6 ; PE$4_2$ 80.3 |
| SM$5_1$ 130.1 ; SM$5_2$ 126.8 | SM$5_1$ 97.8 ; SM$5_2$ 111.9 | PE$5_1$ 115.0 ; PE$5_2$ 108.1 |
| SM$6_1$ 104.4 ; SM$6_2$ 81.9 | SM$6_1$ 87.3 ; SM$6_2$ 132.9 | PE$6_1$ 88.2 ; PE$6_2$ 64.0 |
| SM$7_1$ 90.1 ; SM$7_2$ 95.6 | SM$7_1$ 88.5 ; SM$7_2$ 91.5 | PE$7_1$ 88.4 ; PE$7_2$ 115.7 |

**Figure 4–figure supplement 2.** Average *B*-factors of acyl chains in structures of AQP0 in different lipid bilayers. (**A**) AQP0 in a bilayer formed with a molar sphingomyelin:cholesterol ratio of 1:2. (**B**) AQP0 in a bilayer formed with a molar sphingomyelin:cholesterol ratio of 2:1. (**C**) AQP0 in a bilayer formed with *E. coli* polar lipids. The two acyl chains of each lipid are labeled in the top panel with subscripts and their average *B*-factors are tabulated below.

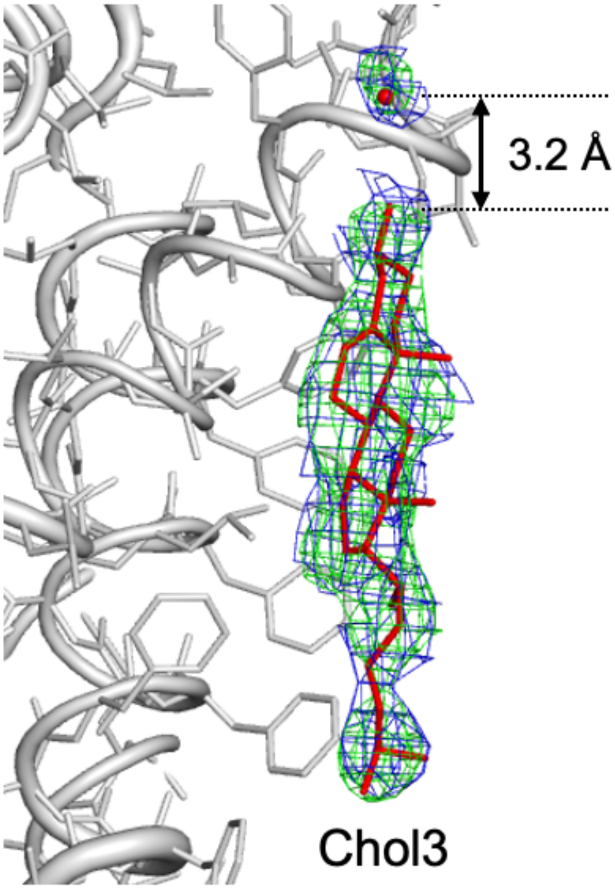


**Figure 4–figure supplement 3.** The hydroxyl head group of Chol3 makes a hydrogen bond with a water molecule. The distance of the water molecule from the Chol3 hydroxyl group is 3.2 Å. AQP0 is shown in gray ribbon and sticks representation, Chol3 is shown as red sticks, and water as a red sphere. The 2Fo-Fc map for Chol3 and the water molecule is shown in green and the composite-omit map in blue.

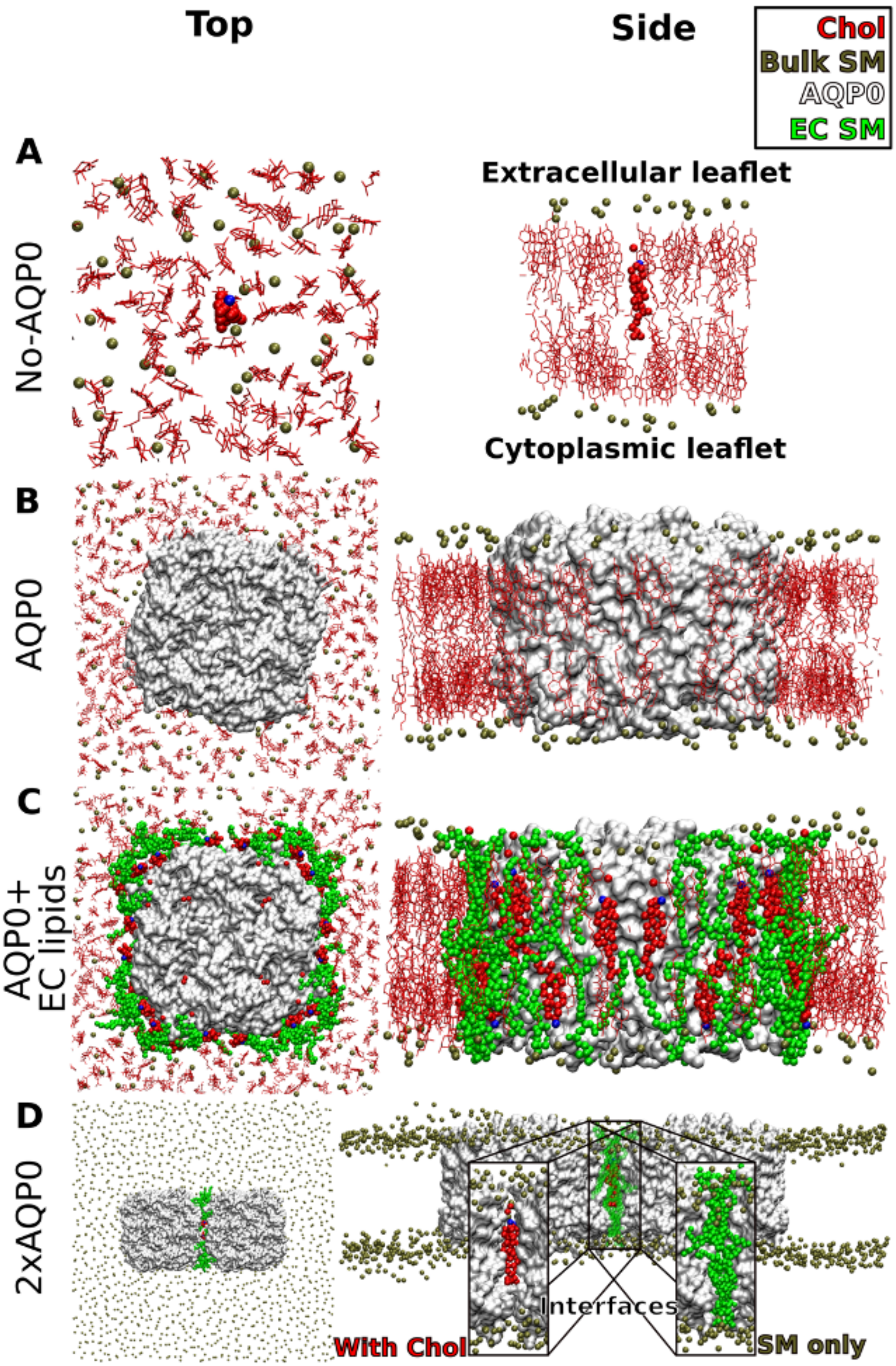


**Figure 5-figure supplement 1.** Molecular dynamics simulations of AQP0 in sphingomyelin/cholesterol membrane systems. (**A-D**) Top views (left panels) and side views (right panels) of the simulated systems. AQP0 tetramers are shown in white surface representation. Lipids in positions seen in the electron crystallographic $AQP0_{1SM:2Chol}$ structure are shown in VDW representation; sphingomyelin (SM) in green and cholesterol (Chol) in red and blue, respectively. Bulk cholesterol molecules are shown in red stick representation, whereas only the phosphorous atoms of bulk SM lipids are shown as dark tan spheres. The membranes were solvated by explicit water molecules (not shown for clarity). (**A**) In the "No AQP0" systems used as control, lipid-only bilayers consisting of molar SM:Chol mixtures of 1:0, 2:1 and 1:2 (shown) were simulated. In each system, one cholesterol molecule was inserted in the position of the deep cholesterol seen in the $AQP0_{1SM:2Chol}$ structure. (**B**) In the "AQP0" system, a single AQP0 tetramer by itself without the annular lipids was inserted into membranes consisting of molar SM:Chol mixtures of 2:1 and 1:2 (shown). (**C**) In the "AQP0 + EC lipids" system, a single AQP0 tetramer with the lipids observed in the $AQP0_{1SM:2Chol}$ structure was inserted into membranes consisting of molar SM:Chol mixtures of 1:0, 2:1 and 1:2 (shown). Note the deep cholesterols at the center. (**D**) In the "2×AQP0" system, a pair of AQP0 tetramers was inserted into a pure SM membrane. The interfacial lipids between the pair of tetramers were either pure SM (inset "SM only") from the $AQP0_{2SM:1Chol}$ structure or a hybrid interface that replaces the two central SM molecules with the EC deep cholesterol molecules seen in the $AQP0_{1SM:2Chol}$ structure (shown, inset "with Chol").

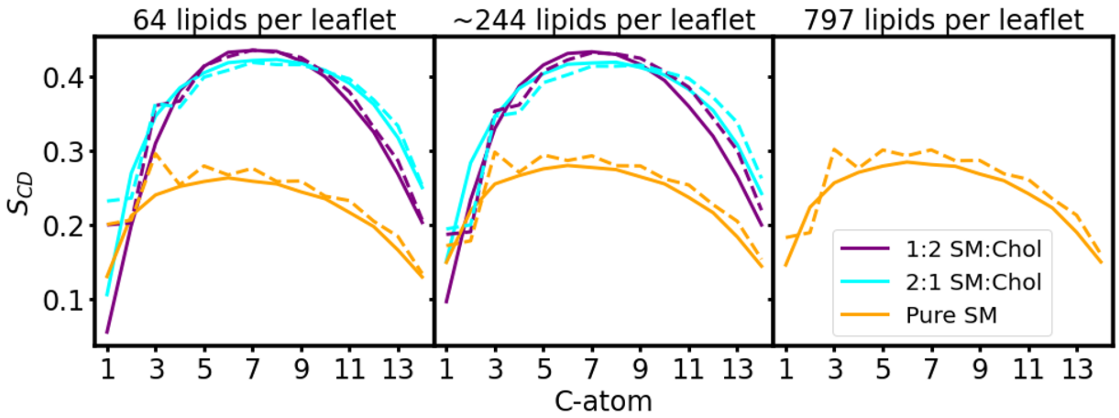


**Figure 5-figure supplement 2.** Deuterium order parameters for the membrane patches in equilibration. Deuterium order parameters $S_{CD}$ for the hydrophobic acyl chains for the SM lipids in a pure SM bilayer (Pure SM, orange), and in SM:Chol membranes at molar mixing ratios of 2:1 (2:1 SM:Chol, cyan) and 1:2 (1:2 SM:Chol, purple). The continuous line shows the values for the SN1 acyl chains and the dashed line shows the values for the SN2 acyl chains. The values correspond to the last 50 ns of simulation, independent of the patch size and total equilibration time. The three boxes correspond to the pure lipid membrane patches, used for the No-AQP0, AQP0 and 2×AQP0 simulations, respectively, before the protein was inserted in the latter two cases. Membrane patch size indicated on top of each box.

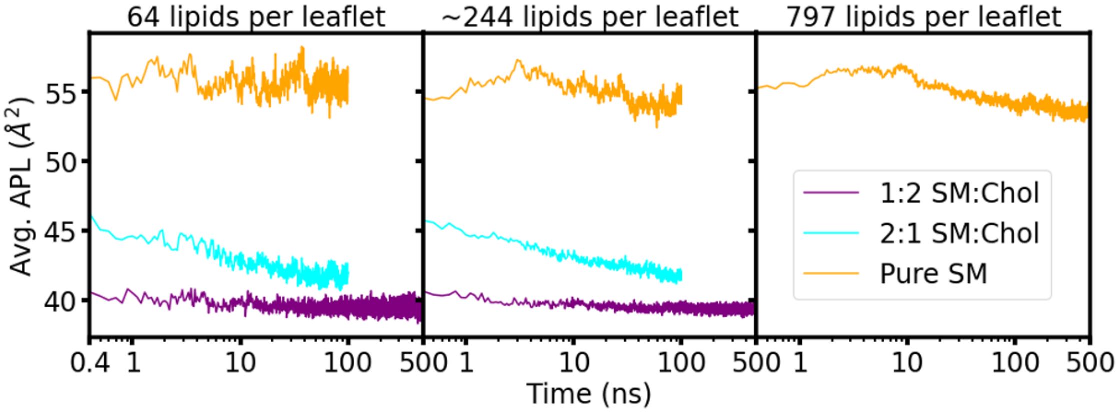


**Figure 5-figure supplement 3.** Average area-per-lipid (APL) for the membrane patches in equilibration. Average APL as a function of time (log scale) is shown for the warm-up NPT simulations after removal of position restraints. The average APL value for the last 50 ns of simulation for the pure SM systems were 55.5±0.7 Å$^2$, 54.2±0.5 Å$^2$ and 53.7±0.2 Å$^2$, for the system with 64, ~244 and 797 lipids per leaflet, respectively.

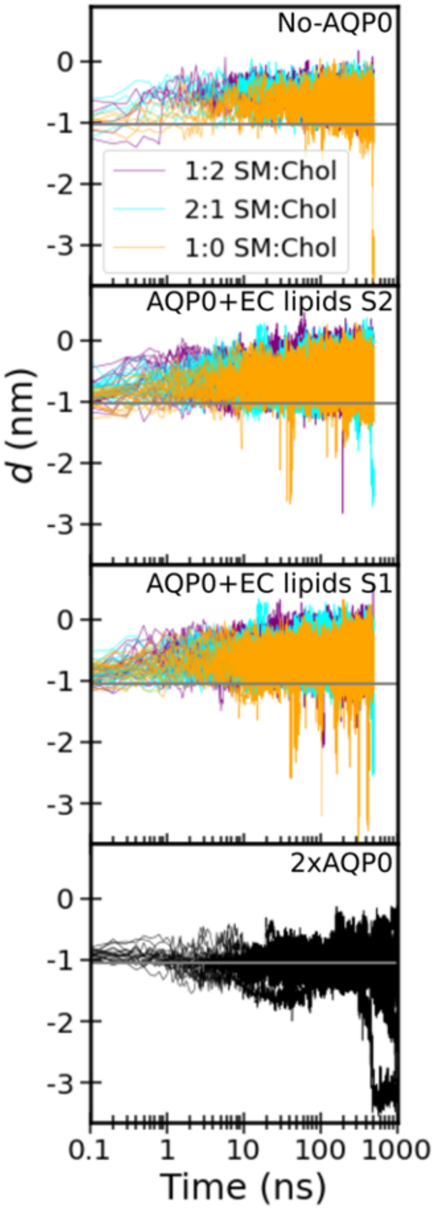


**Figure 6-figure supplement 1.** Timetrace of the insertion depth of cholesterol. Each panel corresponds to one of the four studied systems: “No AQP0”, a pure lipid membrane without AQP0, “AQP0 + EC lipids S2/1”, a membrane with one AQP0 tetramer surrounded by the annular lipids seen in the $AQP0_{1SM:2Chol}$ structure, and “2×AQP0”, a membrane containing a pair of AQP0 tetramers together with the lipids in between them from a hybrid $AQP0_{2SM:1Chol}$ structure that replaces the two central SM molecules with the EC deep cholesterol molecules found in the $AQP0_{1SM:2Chol}$ structure. Color indicates the different SM:Chol ratio and the different curves represent the different replicas for each condition. The horizontal gray line indicates the most probable cholesterol position in the 2×AQP0 system. Distributions shown in main **Figure 6** were obtained from these time traces after discarding the first 100 ns that were considered equilibration time (300 ns for the 2×AQP0 system).

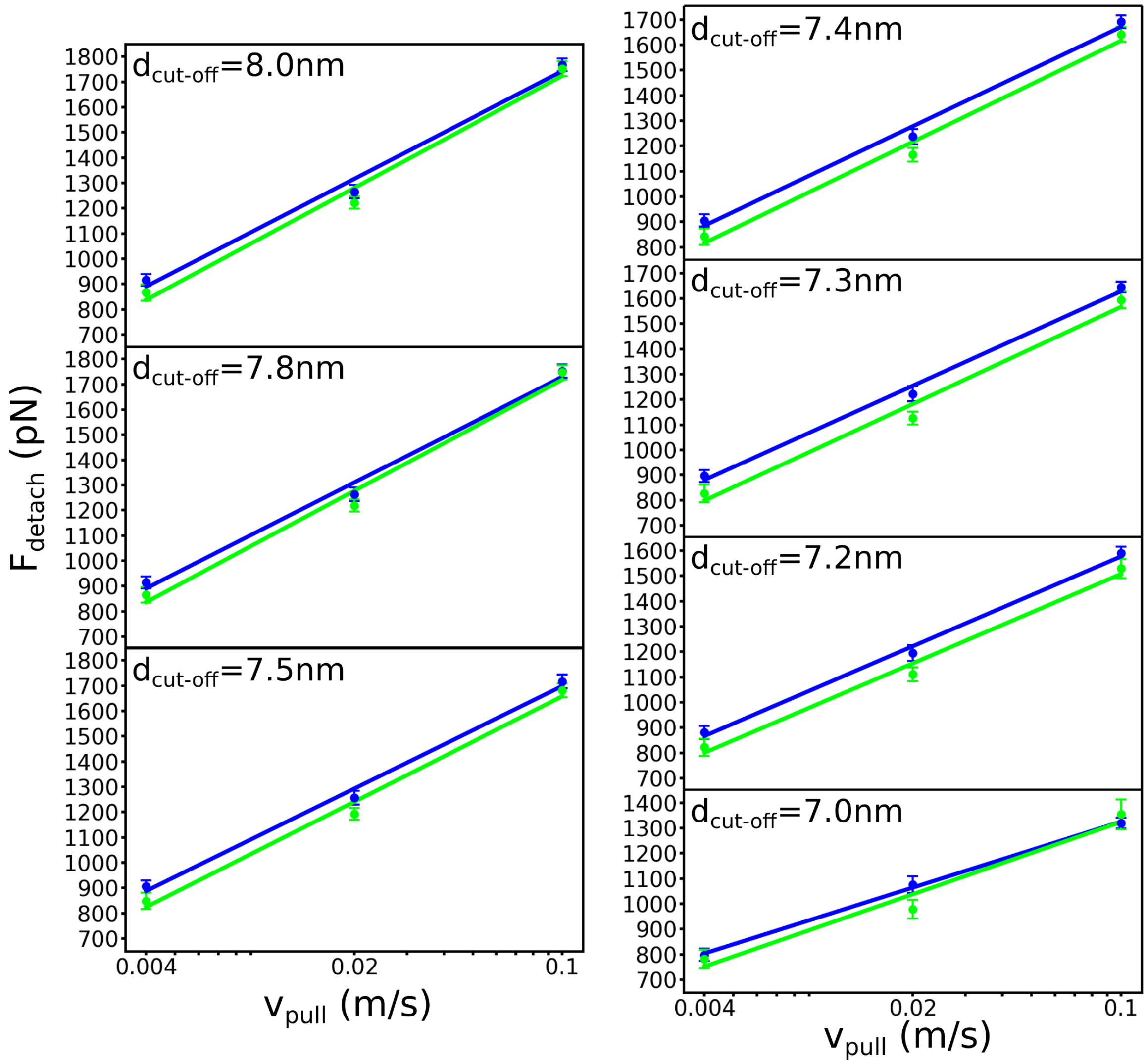


**Figure 7-figure supplement 1.** Detachment force versus pulling rate for different $d_{cut\text{-}off}$ criteria. The cut-off value chosen to define tetramer separation does not alter the conclusion that the presence of deep cholesterol at the interface (blue) results in a more stable association between the tetramers (i.e., requiring a high detachment force) than when the interface only contains sphingomyelin (green).

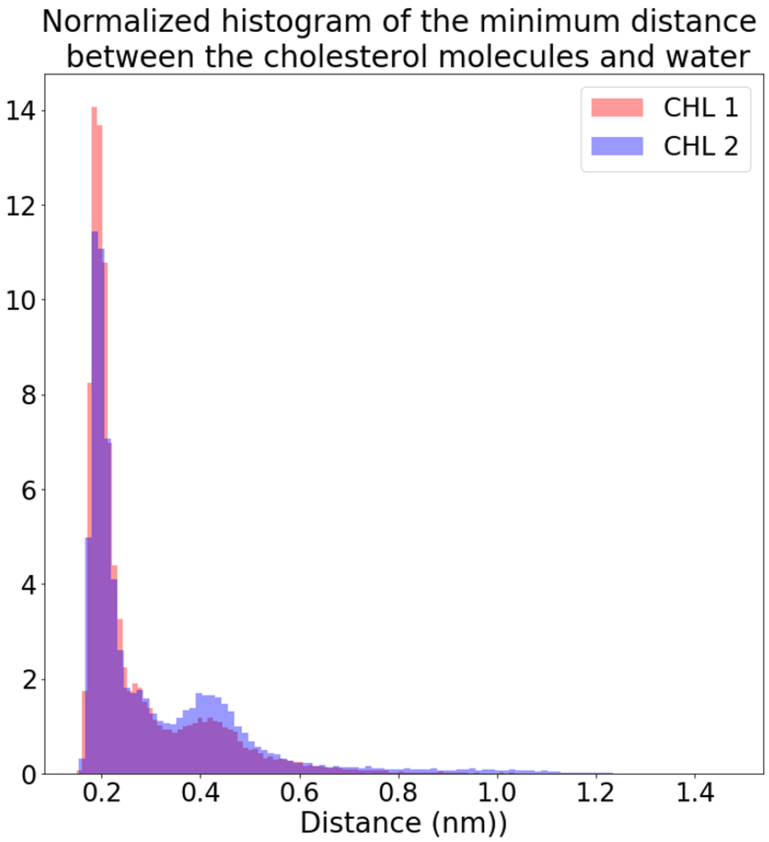


**Figure 8-figure supplement 1.** Distance from each deep cholesterol OH group to the nearest water molecule. Minimum distance between the hydroxyl oxygen atom in each deep cholesterol molecule and the closest water molecule in the 2×AQP0 simulations in equilibrium, shows that they were nearly always in contact.

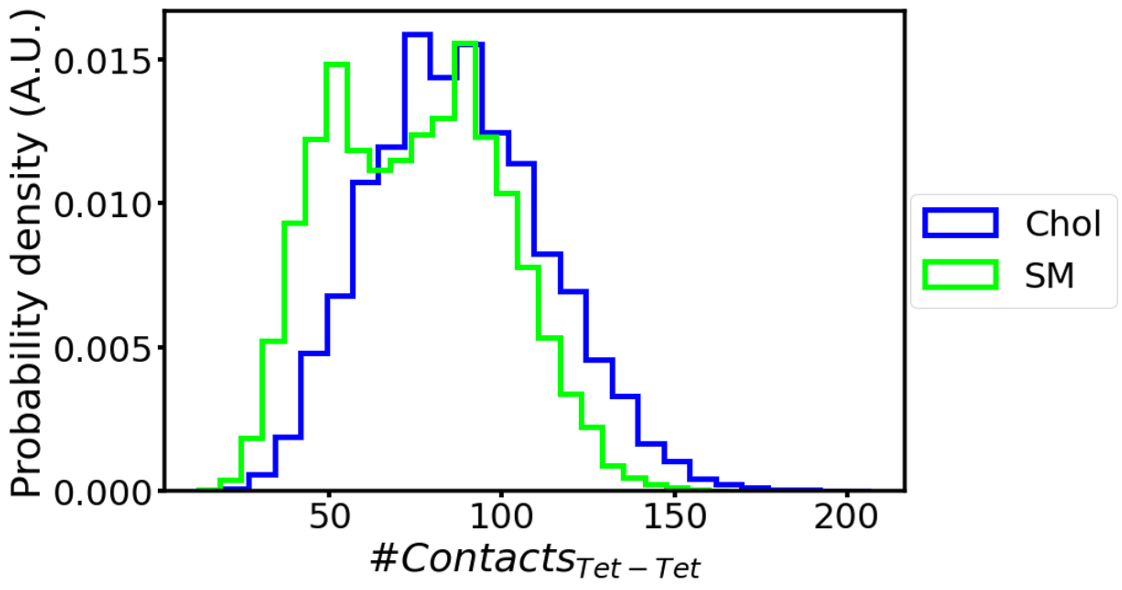


**Figure 8-figure supplement 2.** Contacts between tetramers in 2×AQP0 simulations. Number of contacts between the two tetramers during MD simulations of the 2×AQP0 system in equilibrium for the interface containing only sphingomyelin (SM) and the interface containing deep cholesterols (Chol). The distributions show more protein contacts for the interface with cholesterol in accordance with **Figure 8A-C**.

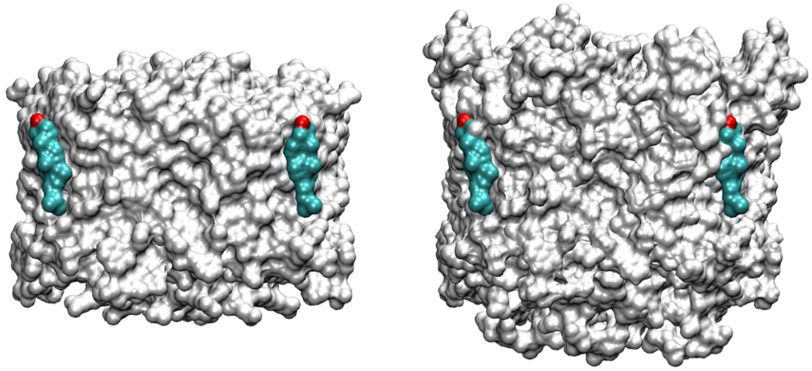


**Figure 8-figure supplement 3.** Surfaces of AQP0 and AQP1. The position of Chol1 seen in the AQP0 structures is indicated at the left and overlapped with the AQP1 structure at the right, which results in steric clashes. The difference in the lateral surfaces of AQP0 and AQP1 may result in a greater association of cholesterol in the Chol1 position in the case of AQP0.
